# The elasto-plastic nano- and microscale compressive behaviour of rehydrated mineralised collagen fibres

**DOI:** 10.1101/2022.09.26.509461

**Authors:** Alexander Groetsch, Aurélien Gourrier, Daniele Casari, Jakob Schwiedrzik, Jonathan D. Shephard, Johann Michler, Philippe K. Zysset, Uwe Wolfram

## Abstract

The multiscale architectural design of bio-based nanostructured materials such as bone enables them to combine unique structure-mechanical properties that surpass classical engineering materials. In biological tissues, water as one of the main components plays an important role in the mechanical interplay, but its influence has not been quantified at the length scale of a mineralised collagen fibre. Here, we combine *in situ* experiments and a statistical constitutive model to identify the elasto-plastic micro- and nanomechanical fibre behaviour under rehydrated conditions. Micropillar compression and simultaneous synchrotron small angle X-ray scattering (SAXS) and X-ray diffraction (XRD) were used to quantify the interplay between fibre, mineralised collagen fibrils and mineral nanocrystals. Rehydration led to a 65% to 75% decrease of fibre yield stress and compressive strength, and a 70% decrease of stiffness with a 3x higher effect on stress than strain values. While in good agreement with bone extracellular matrix, the decrease is 1.5-3x higher compared to micro-indentation and macro-compression. Hydration has a higher influence on mineral than fibril strain while the highest difference to the macroscale was observed comparing mineral and tissue levels. Results suggest that the effect of hydration is strongly mediated by ultrastructural interfaces while corroborating the previously reported water-mediated structuring of bone apatite providing insights towards the mechanical consequences. Results show that the missing reinforcing capacity of surrounding tissue is more pronounced in wet than dry conditions when testing an excised array of fibrils, mainly related to the swelling of fibrils in the matrix. Differences leading to higher compressive strength between mineralised tissues do not seem to depend on the rehydration state while fibril mobilisation follows a similar regime in wet and dry conditions. The lack of kink bands point towards the role of water as an elastic embedding, thus, adapting the way energy is absorbed.

**Statement of significance:** Characterising structure-property-function relationships of biomaterials helps us to elucidate the underlying mechanisms that enables the unique properties of these architectured materials. Experimental and computational methods can advance our understanding towards their complex behaviour providing invaluable insights towards bio-inspired material development. In our study, we present a novel method for biomaterials characterisation. We close a gap of knowledge at the micro- and nanometre length scale by combining synchrotron experiments and a statistical model to describe the behaviour of a rehydrated single mineralised collagen fibre. Results suggest a high influence of hydration on structural interfaces, and the role of water as an elastic embedding. Using a statistical model, we are able to deduce the differences in wet and dry elasto-plastic properties of fibrils and fibres close to their natural hydration state.

## 1 Introduction

The intriguing architecture of bone determines its remarkable ability to combine mechanical properties that are commonly mutually exclusive. While being a light-weight material, it combines high toughness and high strength [1–4] via an elaborate design of its organic and inorganic constituents [5, 6]. Studying bone on its multiple length scales enables us to understand the underlying deformation mechanisms. For bone-related diseases this detailed knowledge is essential to elucidate the related changes in bone’s architecture, especially for pathologies such as osteogenesis imperfecta (OI) where the increased brittleness is related to changes at the molecular level [7]. Thus, not only the information on the bone quantity but also on the micro- and ultrastructure is important. For bio-inspired materials deciphering the structure-mechanical function relationships can inspire new designs including nanostructured metamaterials while identifying potential candidate materials for bone implants [6, 8–12]. In digital healthcare, computational models have the potential to improve diagnosis and treatment strategies [13–16]. They can be used to simulate bone’s mechanical behaviour under multiple loading conditions and to predict its failure behaviour. Such models, however, critically depend on the underlying material description on bone’s multiple length scales [17–21].

Since bone is a multiscale material, different mechanical properties and governing deformation mechanisms are present at its different architectural levels [5, 6, 9, 22–24]. This includes scale transitions with superior strength at the bone extracellular matrix level compared to the macroscopic tissue level [25, 26]. While there is substantial information at the macroscale, there is limited knowledge on ultimate properties such as strength and ultimate strain at the micro- and nanometre lengthscale that might explain these scale transitions [27]. This is in part due to the challenges imposed on *in situ* experiments to extract micro- and nanoscale structure-property relationships of bone’s main constituents, collagen, mineral, and water. At the micro- and nanoscale, bone’s elementary mechanical units are the mineralised collagen fibre and mineralised collagen fibril [9, 23, 28, 29]. A mineralised collagen fibre represents an array of mineralised collagen fibrils [5, 6, 23, 30] which consists of type I collagen molecules and mineral particles of primarily carbonated hydroxyapatite [31, 32]. The fibrils are embedded in an extrafibrillar matrix that consists of a layer of non-collagenous proteins, proteoglycans, extrafibrillar mineral and water [33–37] while sacrificial bonds, cross-links and dilatational bands represent important toughening mechanisms along the scales [36, 38–41]. The collagen molecules within the mineralised collagen fibrils show a staggered arrangement. The intrafibrillar mineral particles are located within the gap zones of these molecules [31, 32, 42–44].

Water makes up about 14-20 wt% of bone [45, 46]. It is reported that the hydration state has an influence on the mechanical properties at the material’s different hierarchical levels as seen in experiments and simulations [47–51]. At the extracellular matrix level, yield stress and compressive strength are reduced by 60% to 75% for rehydrated samples [50]. Bone tissue shows a ductile behaviour at this length scale, also in dry conditions, and a transition to quasi-brittle behaviour at the organ level [25]. For millimetre sized bone samples, hydration leads to a more ductile behaviour [49]. Micropillars from ovine bone extracellular matrix have been tested under quasi-physiologic conditions and confirmed the ductile behaviour on these small length scales [50]. Furthermore, tension and compression of millimetre sized bone samples were combined with SAXS and WAXS/XRD measurements under dry and wet conditions [49, 52, 53]. A reduction of elastic bulk properties was reported as well as a decrease in the ratios between fibril, mineral and macroscopic strains. Water can be found in different regions of the bone matrix influencing the mineral-organic matrix interactions [46, 54]. Apart from pore spaces such as lacunae, canaliculi, and Haversian channels, water is present in a bound form in carbonated apatite as well as in the extracellular bone matrix [46, 47, 54] and small gap regions [55]. It is also present as surface water around mineralised collagen fibrils and mineral platelets which is discussed to affect the energy dissipation [46, 54], thus, influencing mechanisms at the extracellular matrix. It is further postulated that dehydration of the nanocomposite bone has different effects on mineral and collagen, which, in turn, leads to changes in the overall tissue behaviour [47]. However, a crucial gap still exits. The micro- and nanomechanical behaviour of a mineralised collagen fibre under rehydrated conditions has not been studied so far.

Consequently, this study aims at (i) performing micro- and nanomechanical testing of rehydrated mineralised collagen fibres, (ii) quantifying the influence of hydration at the level of a mineralised collagen fibre, mineralised collagen fibrils and mineral nanocrystals, and (iii) using an existing statistical constitutive model as an evaluation method to identify hydrated mechanical fibre properties at molecular and fibrillar levels, thus, capturing the material behaviour.

## 2 Materials and methods

Recently, we reported results on the micro- and nanometre lengthscale compressive behaviour of single mineralised collagen fibres [21, 56]. Micropillar compression [25, 57] and synchrotron radiation X-ray scattering and diffraction techniques (SAXS/XRD) [42, 58–65] were used to extract the mechanical behaviour of a mineralised collagen fibre and the interplay with its mechanical components under dry conditions. Combined with ultrastructural data, those findings were used to develop a statistical elasto-plastic model that explains the micro- and nanoscale fibre behaviour [21]. The extension towards a setup with rehydrated samples now lets us deduce the fibre mechanical properties close to their hydration state in a combined experimental and computational approach.

### 2.1 SAXS/XRD with micropillar compression under rehydrated conditions

Dissection, ultra-milling, laser machining and focused ion beam milling (FIB) were used to separate micropillars from individual mineralised collagen fibres for mechanical testing using a previously developed protocol [56]. Briefly, dissection methods using a scalpel and a diamond band saw (Exakt, Norderstedt, Reichert-Jung) were used to separate a naturally highly mineralised tendon piece from turkey legs (*tarsometatarsus*), an established model system for bone due to its uniaxial fibre arrangement and similar structural set-up at the mineralised collagen fibre level [17, 34, 42, 43, 66–69]. Samples were kept frozen at −22°C until preparation. Before dissection, samples were thawed covered in phosphate buffered saline solution (PBSS). Tendon pieces of 1.5 mm in diameter and 10.0 mm in length were glued into cylindrical aluminium sample holders using a 2-component epoxy resin adhesive (Schnellfest, UHU, Germany). The free end was polished with an ultramiller (Polycut E, Reichert-Jung, Germany). Ultra-short (picosecond) pulsed laser ablation (TruMicro 5250-3C, Trumpf, Germany) was used to cut pre-pillars of 32.85 ± 0.91 μm, centred around a single mineralised collagen fibre (Figure 1). The plasma mediated ablation process minimised the thermal impact during manufacturing. Raman measurements and finite element analyses were used to confirm that the physio-chemical properties were not changed by the laser ablation, and that the ablation process did not exceed the denaturation temperature for dry collagen [56, 70]. Focused ion beam milling (Quanta 3D FEG, FEI, USA) was then used to cut the final micropillars. Micropillars were 6.4 ± 0.6 μm in diameter and 2.05 ± 0.09 in aspect ratio. The possible effects of FIB induced Gallium implantation was assessed by means of Monte Carlo simulations using the software SRIM [71]. Affected layers were shown to be thinner than 30 nm and can, thus, be neglected compared to the micropillar mean diameter with respect to its influence on the mechanical behaviour [56, 72, 73]. To test micropillars from single mineralised collagen fibres in combination with SAXS/XRD under quasi-physiological conditions, a rehydration set-up was developed that allowed it to perform in *situ* micropillar compression experiments at a synchrotron beamline (Figure 2). Before mechanical testing, samples were rehydrated for 2 hours in Hank’s balanced salt solution (HBSS). The state of rehydration was checked using an optical microscope and if necessary, HBSS was added using a pipette. For the alignment between micropillar, flat punch and X-ray beam, excessive HBSS was removed with a pipette so that the micropillar was visible with the microscope (Figure 2). During alignment, the sample was kept rehydrated via diffusion from the micropillar base by filling the HBSS reservoir (Parafilm encasing) at regular intervals. Before compression started, the rehydration state was checked by an optical microscope based on the level of HBSS (Figure 2, bottom-right). We also confirmed that the sample surface was accessible and the flat punch was free of HBSS during the tip approach. The sufficient hydration state was further checked in preliminary tests. These tests showed that 0.05 ml drops of water are not evaporated after 15 min at ambient temperature which is sufficient for the testing at the beamline. In addition, an active diffusion was estimated by applying a 1D diffusion calculation based on Darcy’s law [74] and the small sample dimensions.

**Figure 1:**
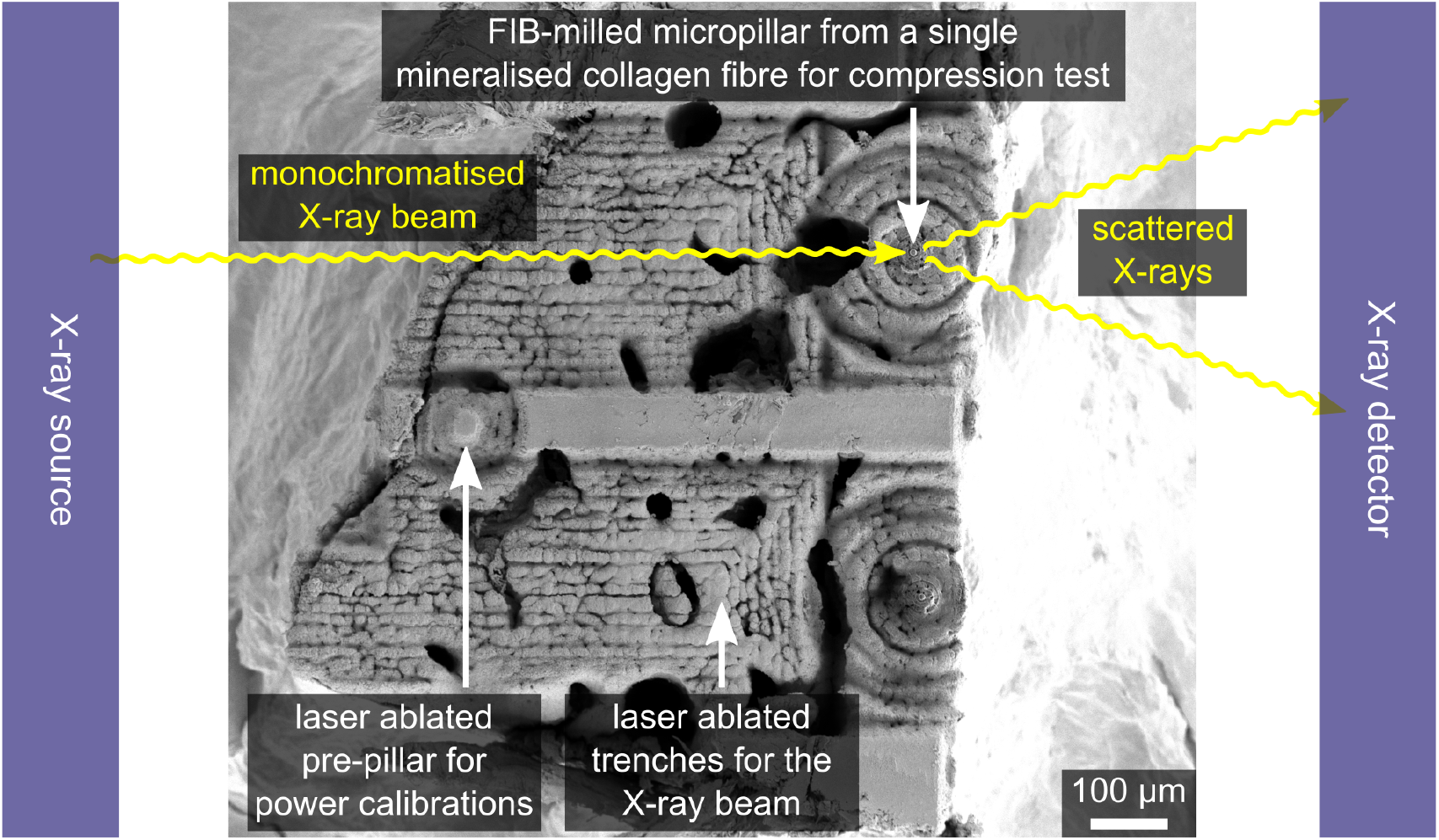
Sample geometry for simultaneous micropillar compression and SAXS/XRD. The SEM image shows the top view of laser ablated trenches to allow the X-ray beam to pass through to FIB-milled micropillars, which were extracted from individual mineralised collagen fibres. They were compressed while SAXS or XRD patterns were taken in a synchrotron setup to quantify the fibril and mineral strain while the apparent fibre behaviour was measured by an integrated microindenter (Figure 2). To account for different tissue reactions during laser processing (natural variability), a pre-test pre-pillar was ablated, checked with focus variation microscopy, and the laser power adjusted if necessary.

**Figure 2:**
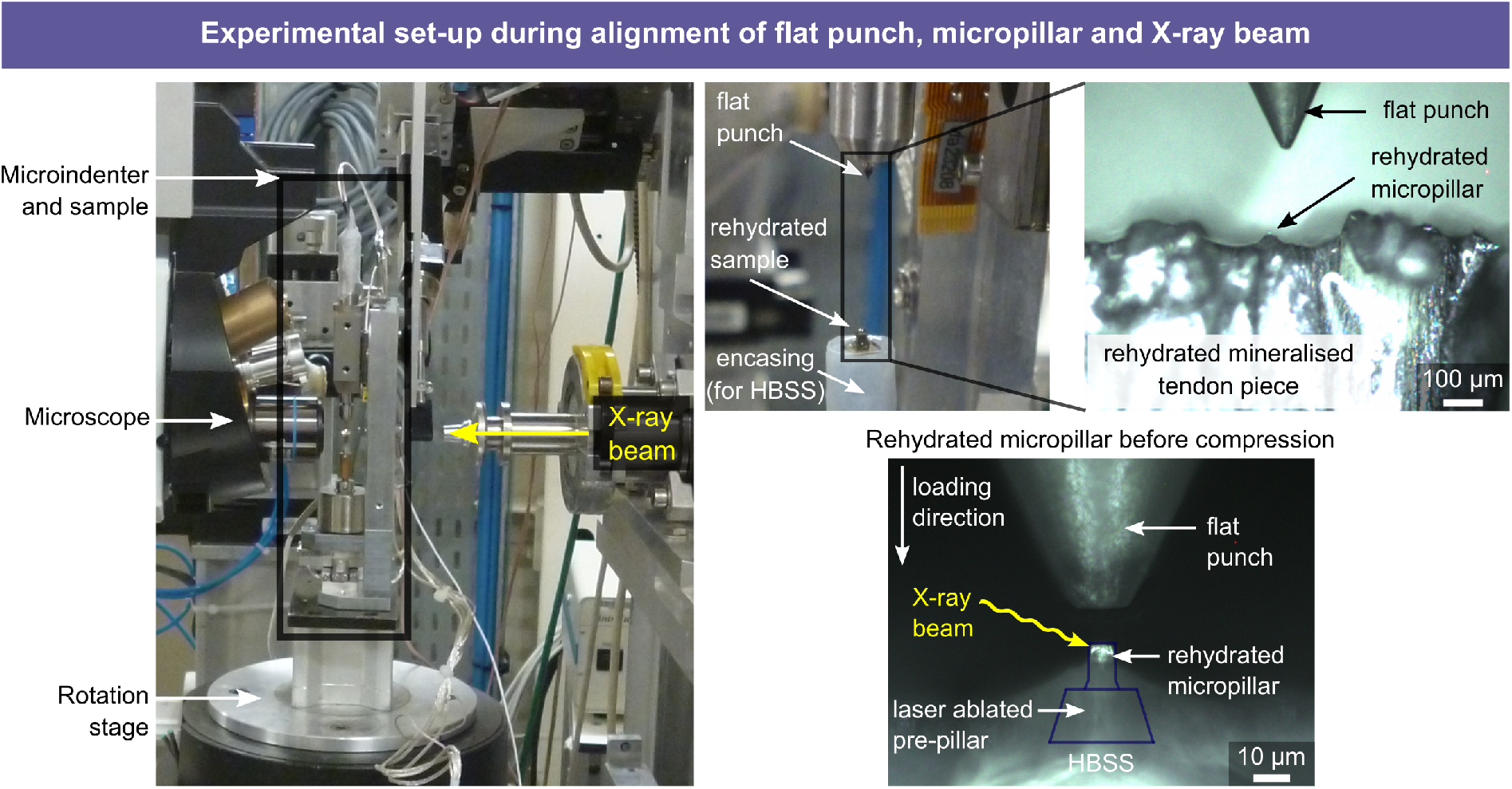
Experimental set-up for the simultaneous micropillar compression and SAXS/XRD measurements of mineralised collagen fibres under rehydrated conditions. The microindenter was implemented into the X-ray set-up of the micro-focus beamline ID13 (ESRF) and placed on a rotation z-stage for the alignment between flat punch, micropillar and X-ray beam. An encasing for the reservoir of Hank’s balanced salt solution (HBSS) was created by wrapping a paraffin-polyethylene foil (Parafilm) around the aluminium tube with the mineralised tendon piece. This reservoir which was refilled at distinct intervals over a period of 2h before testing. The sample was kept rehydrated from the base during the test via diffusion. Bottom-right: The obscured sample outline is indicated and includes the laser ablated pre-pillar. HBSS rose at the micropillar due to surface tension. See [56] for details on the experimental setup under dry conditions.

Mechanical testing and X-ray diffraction followed protocols used for dry tests [56]. Uniaxial micropillar compression experiments were performed with a custom-built microindenter (Alemnis AG, Switzerland) inside a small angle X-ray scattering (SAXS) and X-ray diffraction (XRD) set-up at the ID13 of the European Synchrotron Radiation Facility (ESRF). Micropillars were compressed until failure while being exposed to X-rays at 120 discrete time points. A quasi-static loading protocol at 5 nm/s was used until a mean maximum strain of 12% including partial unloading steps of 50 nm every 150 nm. Either SAXS- or XRD-patterns were recorded during the micropillar compression test to quantify strains of the mineralised collagen fibrils (calcified collagen phase) or mineral particles (mineral phase). X-ray exposure was every 5 s at an energy of E = 13.3 keV. Exposure time for SAXS acquisitions were 75 ms, for XRD 185 ms. The shutter was closed between acquisitions. The beamsize was 5.5 μm in vertical and 7.0 μm in horizontal direction. A single-photon counting detector was used to capture the diffracted/scattered X-ray signal (Eiger, Dectris, Switzerland).

### 2.2 SAXS/XRD and mechanical data analysis

Data analysis for the fibre properties was done in Python (Python Software Foundation, Python Language Reference, version 2.7) [75] and R [76]. Assuming a negligible volume change [77] and a uniaxial stress state within the micropillar [78], engineering stress *σ_eng_* was calculated by dividing the force by the top surface micropillar area. Engineering strain *ε_eng_* was calculated using the displacement and the micropillar height. Geometric measurements were taken with an SEM (Quanta 3D FEG, FEI, USA). Displacement data were frame-compliance and base-compliance corrected. For the base-compliance correction, a modified Sneddon approach [79, 80] was used to account for the elastic sink-in of the micropillar into the substrate beneath (elastic half-space) while considering a fillet radius at the bottom of the micropillar. Engineering stress-strain data were then converted to true stresses and strains via *σ*_33_ = *σ*_eng_(1 + *ε_eng_*) and ln(U_33_) = ln(1 + *ε_eng_*) following previously reported procedures [25, 56, 78] (Figure 3, bottom right). An envelope was fitted to the stress strain curves using a cubic spline with the inflection points of the loading and unloading segments as nodes. This envelope was used to determine the apparent mechanical properties at the mineralised collagen fibre level. Fibre yield values were extracted based on the 0.2% offset criterion, compressive strength and ultimate strain at the maximum of the envelope. Unloading moduli were calculated via linear regression of the unloading segments and their reduction served to estimate damage. The last unloading modulus before the yield point was defined as the apparent Young’s modulus (Figure 3).

**Figure 3:**
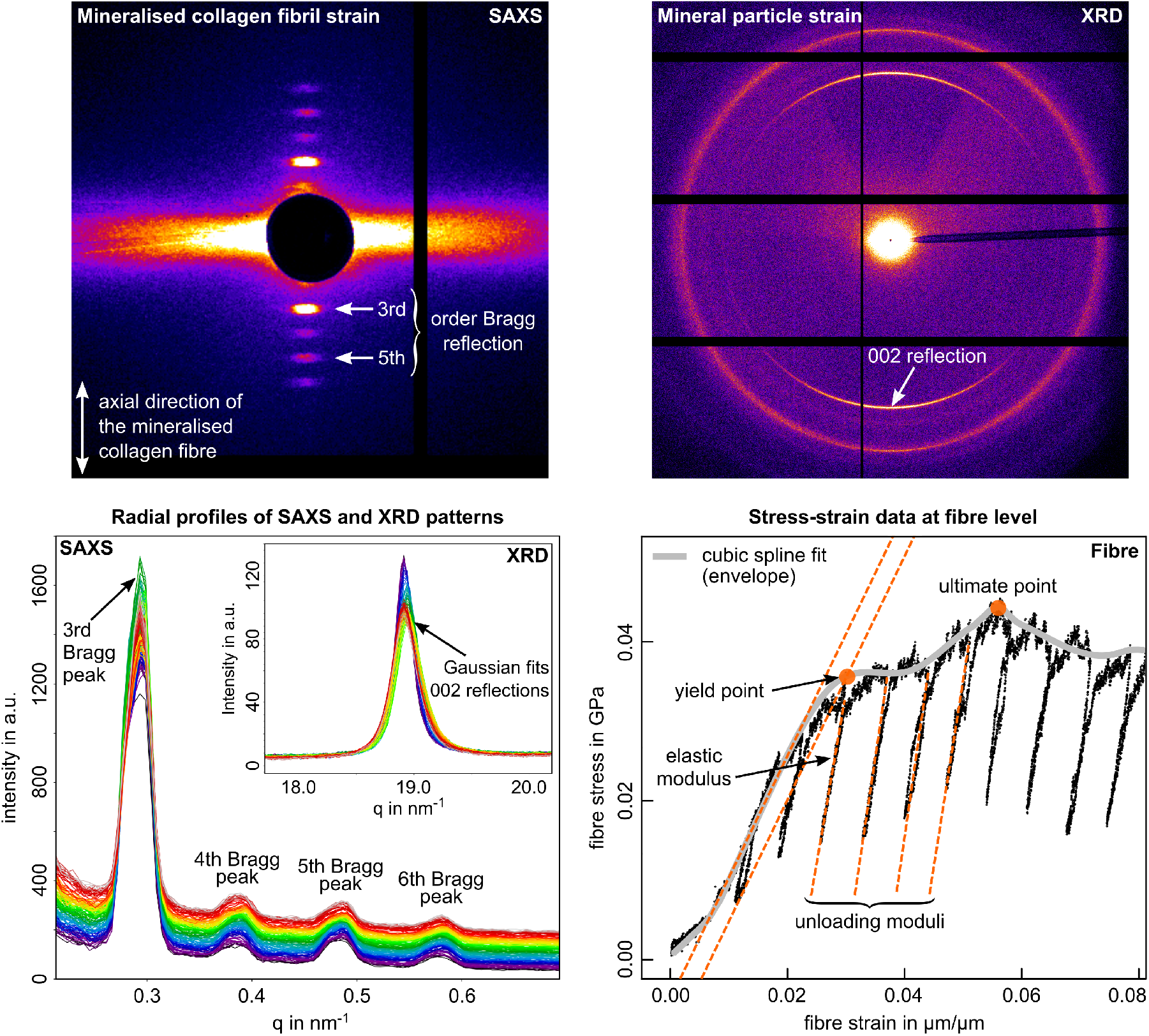
SAXS and XRD patterns as well as data analysis at mineral-, fibril- and fibre-levels. Top-left: SAXS reflections where the 3rd and 5th Bragg order peak were used to determine the mineral collagen fibril strain; Top-right: XRD reflections where the 002 Bragg peak was used to determine the mineral particle strain; Bottom-left: Radial profiles from SAXS and XRD reflections to analyse changes in the position of the wave vector *q* resulting from compressing the fibre micropillar. Peak positions were identified via a Gaussian fit for every of the 120 X-ray acquisitions points per sample. The background (diffuse and parasitic scattering) was fitted with a first order polynomial. The resulting wave vector values *q* were then converted to *d*-values via the relation *q* = 2*π/d* A detailed description of the X-ray data analysis can be found in [56]; Bottom-right: Apparent mechanical properties at the mineralised collagen fibre level were determined from apparent stress-strain curves. A cubic spline (envelope) was fitted to the data using the inflection points as nods. This envelope was used to identify yield values based on the 0.2% offset criterion and ultimate values at the envelope’s maximum. Unloading moduli represent the apparent fibre moduli with the last before the yield point defined as the apparent Young’s modulus.

The deformation of mineralised collagen fibrils and mineral particles can be assessed by analysing changes of the SAXS- or XRD-patterns as a result of the compression. For the SAXS pattern, changes in the Bragg-peaks are related to the changes in the axial arrangement of the intrafibrillar collagen molecules (d period spacing), partly filled with mineral nanocrystals [56, 61]. For the XRD pattern, deformation of the mineral nanocrystals is quantified by tracking changes of the 002 reflection representing changes of a crystal plane perpendicular to the particle c-axis and, thus, to its longitudinal direction. Both the SAXS meridional reflections and the c-axis of the mineral nanocrystals are aligned along the longitudinal direction of the mineralised collagen fibre (Figure 3, top). This allows it to directly compare strain values at fibre-, fibril- and mineral-levels when micropillar compression is combined with SAXS and XRD measurements. Calibration measurements to determine the beam centre, sample-to-detector distance and tilt angle were done using silver behanate (SAXS) and di-aluminium dioxide (XRD). Data analysis for the SAXS and XRD patterns was done with a Python based custom-written software called ’pySXIm’ (courtesy of Aurélien Gourrier, Université Grenoble Alpes, LIPhy, France; converted for Linux by Alexander Groetsch), and the ESRF-developed software Fit2d [81]. Both PCs for the mechanical testing and X-ray acquisition were integrated into the same computer network as to synchronise the datasets for the analysis based on the time stamps in the data log files.

For the direct comparison between fibre, fibril and mineral strain, strain data were synchronised based on time stamps in the data log files for both micropillar compression and SAXS/XRD measurements. In total, 120 acquisition points were accessible throughout the compressive loading and data subsets from the fibre-, fibril- and mineral levels were identified for the strain ratio calculations [56]. The overall strain ratios in the elastic region between the fibre, fibril and mineral deformations were determined at the steepest slope of the cubic spline fit in the elastic region of the mineral-vs-fibre-strain and fibril-vs-fibre-strain plots (Figure 8). Elastic regions were identified based on the previous analysis of the apparent fibre mechanical properties (Figure 3).

### 2.3 Constitutive model as a statistical evaluation tool for the biomaterial behaviour

The connection between the apparent mechanical properties at the mineralised collagen fibre level (microscale) and the deformation of the fibre components (nanoscale) was quantified based on comparing the strain of the mineral particles, the mineralised collagen fibrils and the mineralised collagen fibre [21, 56]. We have developed a constitutive model [21] to interpret such experiments and to quantify the compressive behaviour via those strain ratios. The model is necessary to account for experimental artefacts, which then allows us to simulate the micro- and nanomechanical behaviour of a mineralised collagen fibre, and to deduce its mechanical properties. It operates at the continuum-, molecular- and crystalline-levels covering the fibre, fibrils, collagen molecules and mineral nanocrystals. Material properties for the model follow a statistical normal distribution to account for natural heterogeneities in the material behaviour. Two nested shear lag models are used to calculate elastic properties at the fibre- and fibril level while two inelastic strain mechanisms capture the plasticity simulating the two interfaces in the intra- and extrafibrillar phases of mineral-collagen-mineral, and fibril-matrix-fibril. A parallel arrangement of several model elements represent the parallel array of fibrils within a fibre. The model outputs strain ratio distributions between the constitutive phases and the apparent fibre strain to allow a direct comparison to the statistical values from synchrotron experiments as a mean from the illuminated volume. In the model, we included a non-linear gradual recruitment of mineralised collagen fibrils, which was informed by ultrastructural data from synchrotron phase-contrast nanoCT scans [21], and which accounted for an experimental artefact. The high agreement between model and experiment [21] allows us to interpret the test results under rehydrated conditions.

We include hydration by adapting collagen stiffness [47, 82, 83] and the overall experimentally determined apparent yield strain 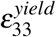 and related parameters (fibril and EFM yield strains) [21] compared to the dry tested micropillars [56]. Thus, a reduced value for the modulus of collagen molecules of 0.9 GPa was used, compared to 1.42 GPa for the dry fibre model [21]. For the formulation of our model, it was further assumed that the fibril volume fraction was kept constant. A full overview of model input parameters are given in Table 5 (Appendix). By combining the knowledge from the experiments and the use of the model we can then derive the actual mechanical properties at the micro- and nanometre lenghtscale including the stiffness of a mineralised collagen fibril. A detailed description of the statistical constitutive model can be found in our previous work [21].

### 2.4 SEM based post-test and failure mode analysis

Failure modes were analysed based on SEM images (E = 2 keV, tilt angle = 45°, Quanta 650 FEG SEM, FEI, USA). Samples were not re-sputtered for post-test failure analyses to avoid that features are being disguised by the gold sputtering. The micropillars are considered as fibril reinforced composites and failure mechanisms were categorised based on the classification scheme previously developed for dry testing [56]. This categorisation was derived from compressive failure modes and toughening mechanisms found in non-biological fibre-reinforced composites and bone tissue where more than one failure mechanism could be present in a single specimen.

### 2.5 Statistical analysis

Statistical analyses were done in R [84] and Python [85, 86]. Quantile-quantile plots and Shapiro-Wilk tests [87] were used to verify that the data were normally distributed and mean ± standard deviation were calculated. In addition, raw data of the samples by means of distribution independent median, minimum and maximum values are presented. To check for statistically significant differences between SAXS and XRD groups, and between the results from rehydrated and dry tests [56], a student t-test was used with a significance level of p=0.05.

## 3 Results

### 3.1 Mechanical properties at mineralised collagen fibre level and influence of hydration

In total, 14 micropillars were fabricated and 10 of the 14 rehydrated samples could be used for the analysis and the comparison to the dry testing. From the four remaining samples, two were used for pre-tests and two were lost due to technical difficulties. Data from rehydrated samples (5 SAXS, 5 XRD) were compared to those tested previously under dry conditions (6 SAXS, 5 XRD) [56]. No significant difference was found for the fibre mechanical properties between the SAXS and XRD group and data were pooled for further analysis. Shapiro-Wilk tests showed that data were normally distributed. Mean values for the experimental fibre mechanical properties are presented in Table 1 for yield stress 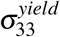, yield strain 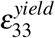, compressive strength 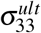, ultimate strain 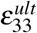 and Young’s modulus *ϵ_fibre_*. In addition, absolute values including median, minimum, and maximum are presented (Figure 4, left). No X-ray beam induced deterioration of the apparent material properties was found based on the unloading moduli (stiffness). No significant increase in damage, estimated as a reduction of the unloading moduli, was detected, up to a plastic strain of 10%. Hydration led to a three times higher reduction in stress than strain values. Due to hydration, a mean reduction of around 75% was observed for the yield stress 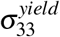 and compressive strength 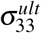. For the yield strain 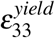 and ultimate strain 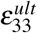, the rehydration led to a reduction of around 25%. The apparent Young’s modulus *ϵ_fibre_* was reduced by 60-70%. All differences between results from the rehydrated and dry testing were significant (Tables 1 and 2).

**Figure 4:**
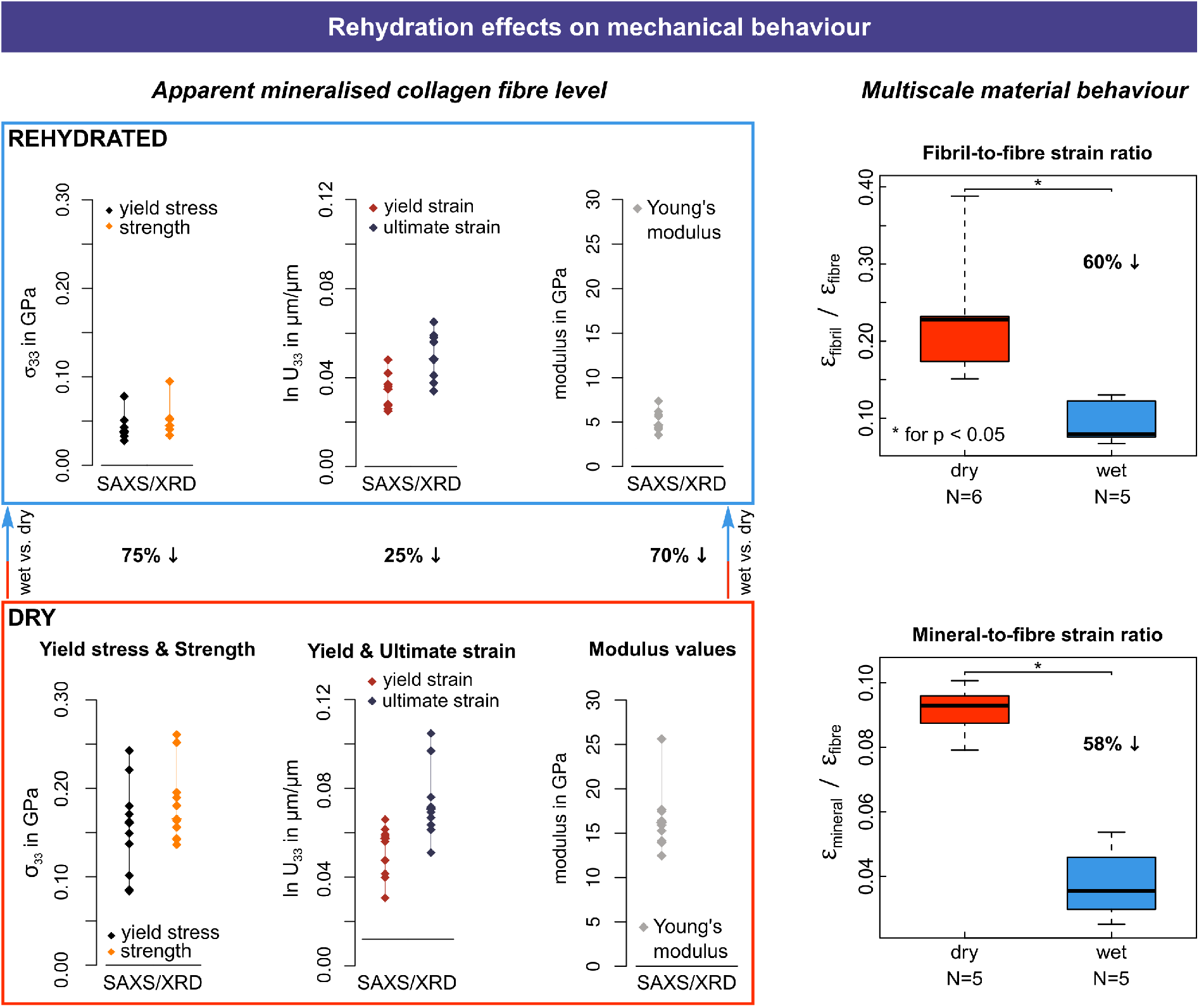
Experimentally determined rehydration effects on the mechanical properties tested under dry (N=11) and rehydrated (N=10) conditions. Left: Effect at the apparent mineralised collagen level with minimum, median and maximum values for the apparent yield point, ultimate point and Young’s modulus. Right: Multiscale strain ratios comparing values at fibre-, fibril- and mineral-levels. Dry testing results from [56]

**Table 1:**
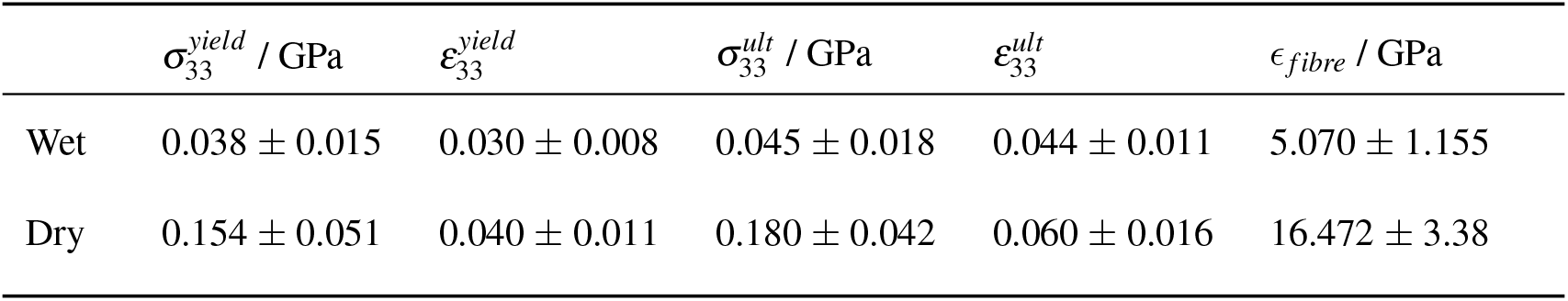
Apparent mechanical properties of a mineralised collagen fibre under rehydrated (N=10) and dry conditions (N=11). Results from dry micropillar compression tests from [56].

**Table 2:** Comparison of rehydrated and dry testing results for the apparent mechanical properties at the mineralised collagen fibre level including p values.

| | $\sigma_{33}^{\text{yield}}$ | $\epsilon_{33}^{\text{yield}}$ | $\sigma_{33}^{\text{ult}}$ | $\epsilon_{33}^{\text{ult}}$ | $\epsilon_{\text{fibre}}$ |
| --- | --- | --- | --- | --- | --- |
| Factor wet/dry compared to dry testing results [56] | 0.247 | 0.744 | 0.251 | 0.741 | 0.308 |
| p value | < 0.00001 | 0.032 | < 0.00001 | 0.014 | < 0.00001 |

### 3.2 Mechanical properties of mineralised collagen fibrils and mineral particles with multiscale strain ratios and influence of hydration

Experiments and simulations show that the mineral strain is smaller compared to the fibril strain and the fibril strain smaller than the apparent fibre strain (Figure 4). Experimentally, this led to ratios in the elastic region of fibre:fibril:mineral of around 53:5:2. Mean values, thus, showed that only around 10% of strain is taken up by the mineralised collagen fibrils compared to the apparent fibre strain, and 4% by the mineral particles (Table 3 and Figure 8). Mineral particle strain was on average 60% smaller than mineralised collagen fibril strain. The effect of hydration was nearly the same for the fibril-to-fibre and the mineral-to-fibre strain ratios with a difference of around 1%. For both the mineralised collagen fibrils and mineral particles, rehydration led to around 60% reduction of strain in the elastic region, with 59.5% for fibril-to-fibre, and 58.4% for mineral-to-fibre (Table 3 and Figure 6). These values are biased by the gradual recruitment of fibrils upon compression (Section 2.3 and [21] (corrected simulation based values are presented in the next paragraph). Rehydrated testing showed relative ratios of 53:5:2 for the fibre, fibril and mineral strain (elastic region) compared to 22:5:2 for the dry testing [56]. Thus, compared to the dry testing, the experimental factor between the mineral and fibril strains remained the same with around 40% less strain taken up by the mineral particles compared to the mineralised collagen fibrils (dry: 0.39, wet: 0.40). Rehydration led to a significant difference of the mean fibril-to-fibre strain ratio from 0.23 to 0.10, and for the mean mineral-to-fibre strain ratio from 0.09 to 0.04, both measured in the elastic region (Section 3.2 and Table 3).

**Table 3:** Experimental ratios (wet only) and factors between fibril and fibre strain as well as mineral and fibre strain under rehydrated and dry testing conditions determined in the elastic region. SAXS (N=5), XRD (N=5)

| | $\epsilon_{33}^{fibril} / \epsilon_{33}^{fibre}$ | $\epsilon_{33}^{mineral} / \epsilon_{33}^{fibre}$ | $\epsilon_{33}^{mineral} / \epsilon_{33}^{fibril}$ |
| --- | --- | --- | --- |
| Strain ratio (wet)<br>(elastic region) | $0.095 \pm 0.029$ | $0.038 \pm 0.012$ | $0.400 \pm 0.021$ |
| Factor wet/dry<br>(compared to dry<br>testing results [56]) | 0.405 | 0.416 | 0.973 |
| p value | 0.004 | < 0.0001 | > 0.05 |

Using the statistical model [21] (Section 2.3) to simulate the mineralised collagen fibre behaviour, we observe a fibre yield stress 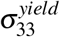 of 0.040 GPa, a yield strain 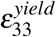 of 0.029, a compressive strength of 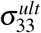 of 0.044 GPa, and an apparent fibre modulus *ϵ_fibre_* of 7.026 GPa (Figure 5). With the shear lag model [21] (Section 2.3) we obtain a mineralised collagen fibril stiffness *ϵ_fibril_* of 15.921 GPa. Solving the experimental artefact using the fibril recruitment model (Figures 5 and 6), mean values for the strain ratios at the yield point were 0.093 between fibrils and fibre as well as 0.035 between mineral nanocrystals and fibre. The ratio between mineral nanocrystals and fibrils was 0.376. Comparing fibre stiffness, yield stress, compressive strength and strain ratios between fibre, fibril and mineral, we observe an agreement of 88.8 ± 8.3 % between simulation results and experiments. We can use the model to exclude the gradual fibril recruitment and derive the actual fibre behaviour. By doing so, we observe a mean fibril-to-fibre strain ratio of around 0.41, a mean mineral-to-fibre strain ratio of 0.13, and a mean mineral-to-fibril strain ratio of 0.32 at the moment the fibre yields (Figure 6). Under dry conditions, corresponding values are 0.70, 0.37 and 0.53, which led to ratios betweem wet and dry of 0.59, 0.35 and 0.60. Furthermore, a 12% higher hydration influence on the mineral than fibril strain was observed leading to a higher reduction in strain in the mineral phase (Table 4).

**Figure 5:**
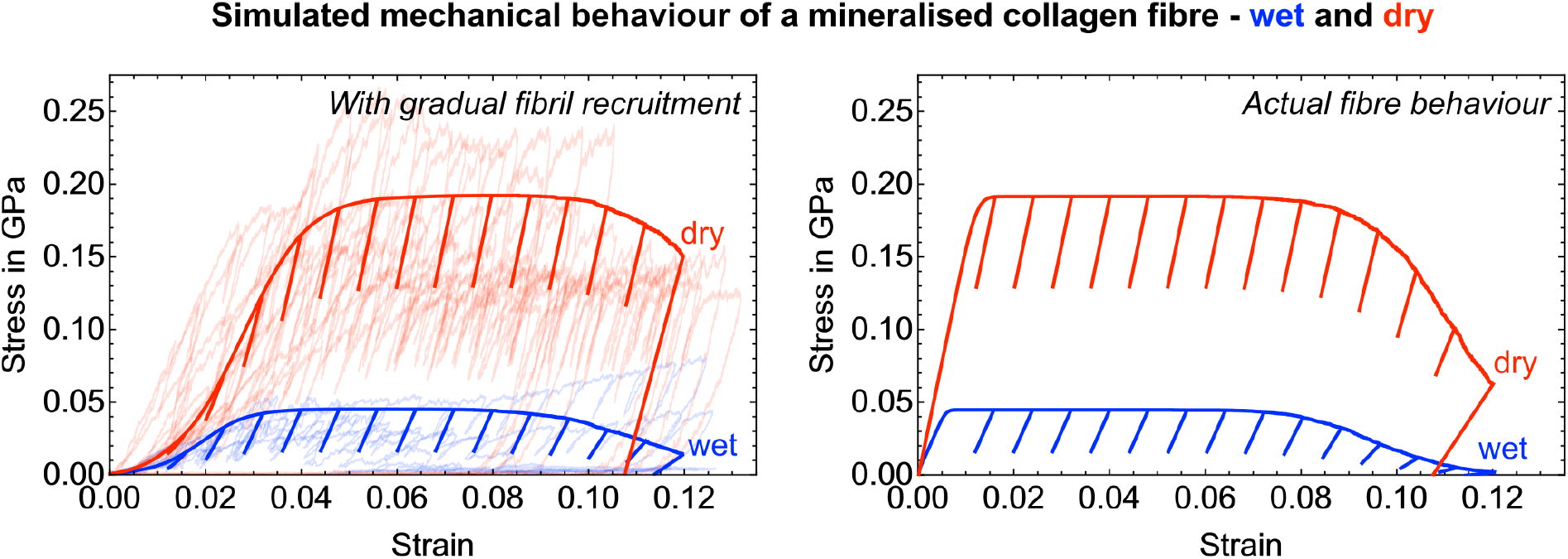
Model outcome at mineralised collagen fibre level for both wet and dry conditions. Left: With gradual fibril recruitment (see Section 2.3 and [21]). Experimental curves are shown in the background in faded red (from [56]) and blue. Right: Actual fibre behaviour.

**Figure 6:**
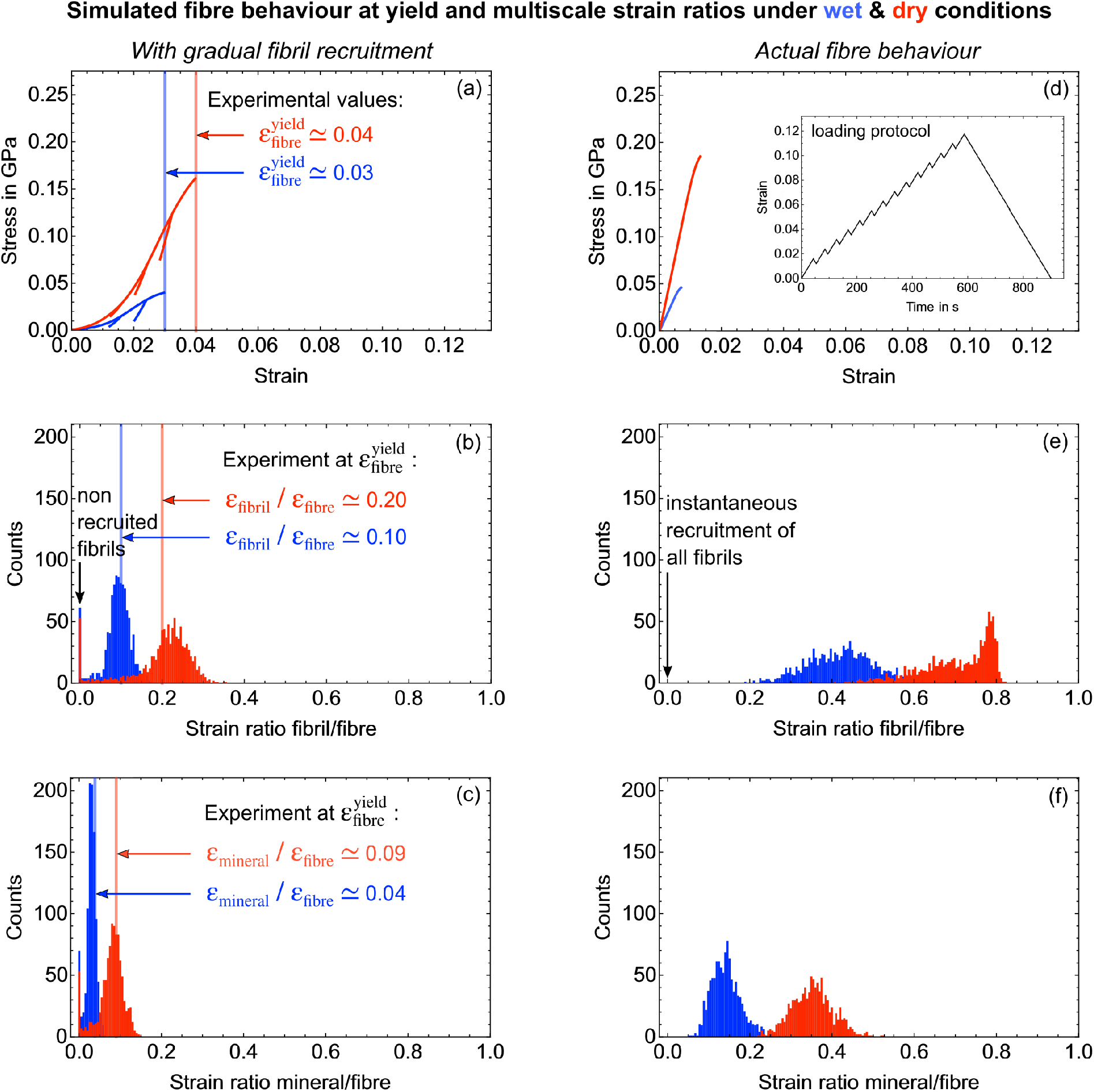
Model outcome for the multiscale mineralised collagen fibre behaviour as simulated at the numerical yield point for both wet and dry conditions. The top row shows the fibre stress-strain data, the second and third rows the strain ratio distributions between fibril and fibre and mineral and fibre levels. Left (a-c): Data when accounting for a gradual fibril recruitment (experimental artefact). Right (d-f): Data where the recruitment is excluded, thus depicting the actual micro- and nanomechanical behaviour of a hydrated mineralised collagen fibre. The inlay in (d) shows the loading protocol for the strain driven simulation.

**Table 4:** Simulated ratios and factors between fibril and fibre strain as well as mineral and fibre strain under rehydrated and dry testing conditions determined at the numerical yield point.

| | $\epsilon_{33}^{fibril} / \epsilon_{33}^{fibre}$ | $\epsilon_{33}^{mineral} / \epsilon_{33}^{fibre}$ | $\epsilon_{33}^{mineral} / \epsilon_{33}^{fibril}$ |
| --- | --- | --- | --- |
| Strain ratio (wet)<br>(at numerical yield) | 0.41 | 0.13 | 0.32 |
| Strain ratio (dry)<br>(at numerical yield) [21] | 0.70 | 0.37 | 0.53 |
| Factor wet/dry | 0.59 | 0.35 | 0.60 |

### 3.3 SEM based failure mode analysis

For the majority of wet tested micropillars, mushrooming at the top surface was observed. In many cases, this was combined with microbuckling or fracture of mineralised collagen fibrils (Figure 7). The occurrence of mushrooming failure modes supports a homogeneous hydration state along the micropillar. If different hydration states would have been present, the lower end of the micropillar would have been more compliant, and strain would localise there leading to failure at the bottom. Comparing failure modes in compressed micropillars under dry conditions [56] fibril-matrix interface failure, axial splitting, fibril microbuckling and fibril fracture occurred in rehydrated and dry tested samples. Bulging and kink bands were only observed in dry tested micropillars, mushrooming only in rehydrated samples.

**Figure 7:**
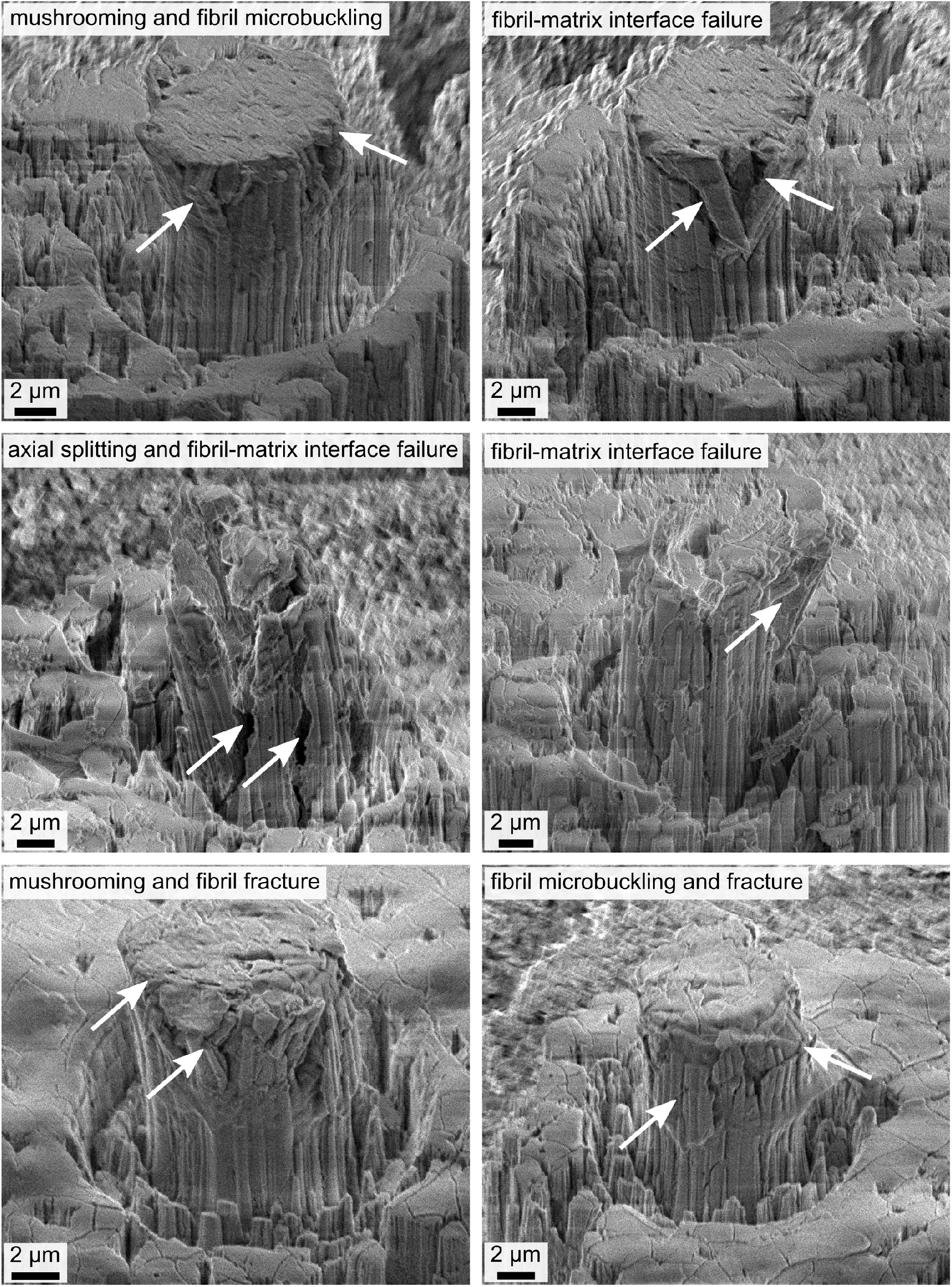
Failure modes of rehydrated micropillars tested under compression. Several failure modes may combine in any one specimen. For the majority of wet tested micropillars, mushrooming at the top surface was observed. In many cases, this was combined with microbuckling or fracture of mineralised collagen fibrils.

## 4 Discussion

We used a combination of multiscale experiment and statistical constitutive model to capture the micro- and nanomechanical behaviour of a mineralised collagen fibre under quasi-physiologic conditions. This included its apparent behaviour as well as the interplay with its components, mineralised collagen fibrils and mineral nanocrystals. To the best of our knowledge this is the first report on a combined experimental setup that integrates micropillar compression of rehydrated single mineralised collagen fibres with synchrotron small angle X-ray scattering (SAXS) and X-ray diffraction (XRD). Using our model as a statistical evaluation method [21] allowed it to account for experimental artefacts aiming to deduce the actual structure-mechanical properties of a single mineralised collagen fibre, and the dedicated effect that water has on the different constitutive phases. A direct comparison to experiments under dry conditions allowed us to quantify the effect of hydration on the multiscale material behaviour.

### 4.1 Apparent mechanical behaviour of rehydrated fibres and effect of hydration

Simulation results showed that the gradual fibril recruitment (experimental artefact) has a significant influence on both fibre yield values and the interaction between fibre constituents (Section 4.2 & Figure 6). Our results show that the mean apparent fibre Young’s modulus is around 50% lower than wet microindentation moduli reported for mineralised turkey leg tendon [88]. Differences can be rationalised by the fact that the microindentation imprints were done on an array of mineralised collagen fibres covering the pore spaces in between fibres. The effect of hydration on these porosities, and the corresponding deformation and energy dissipation mechanisms might result in different stiffness values compared to our results. We also have to consider the swelling and deformation of an excised array of fibrils. In contrast to microindentation tests, testing a free micropillar of an array of uniaxially arranged fibrils [21, 56] lacks the support of surrounding fibres. Since our dry testing results [56] compare well with nanoindentation [88], considering this different set of boundary conditions suggests that we observe lubrication and swelling but that this is not blocked by surrounding tissue, thus losing a reinforcing and stabilising element. This was not found in dry samples [56]. Compared to micropillar compression of rehydrated ovine cortical bone extracellular matrix [50], Young’s modulus, yield stress and compressive strength are about four times lower. This can be related to the lower mineralisation that we found in our dry samples of mineralised turkey leg tendon [21] compared to bone [89]. Generalising our statistical constitutive model towards bone extracellular matrix showed that a higher mineralisation and differences in the interfibrillar surface-to-volume ratio [21] can explain the higher compressive strength in dry bone micropillars [25]. Given that the rehydration related decrease in compressive strength is the same for bone [50] and mineralised turkey leg tendon (75%) (as observed in the present study) this effect, however, seems not to depend on the rehydration state of the tissue. This, in turn, also points to the interesting observation that the mobilisation of fibrils follows a similar regime under dry and wet conditions. Regarding the influence of hydration on the elastic fibre behaviour we observed a decrease of stiffness between 60% and 70% compared to dry testing of the same material [56]. Our observed reduction in yield values are in line with the 65% decrease in yield properties reported for micropillar compression tests of ovine bone extracellular matrix [50]. Compared to wet and dry microindentation results, these values are higher where a reduction of 20-40% in the elastic modulus is reported for rehydrated ovine bone [50], 30% for rehydrated trabecular bone [48] and 6% for rehydrated mineralised turkey leg tendon [88]. Compressive tests on mm-sizes human femur samples showed a decrease in the elastic modulus of around 40% and a decrease in the compressive strength of around 55% [49]. Differences to the macroscale point towards the effect of hydration on additional interfaces when testing samples above the microscale. Our results, thus, suggest a strong influence of ultrastructural interfaces on the hydration effect, combined with the missing stabilising influence of surrounding tissue as discussed earlier. The swelling effects are also expected to be smaller at the macroscale due to a smaller surface-to-volume ratio and overall higher heterogeneity of the ultrastructure.

Studying the effect of water on the fibre constituents, we observe that the model identified a higher influence on the mineral than the fibril strain - an effect also shown for mm-sized bone samples in tension [52] and compression [49]. Compared to dry tissue [56], we observe smaller values for both the mineral and fibril strain when the fibres are wet (Figure 6). As discussed for the mineral strain, the lower strain can be explained by a lubrication of the fibrils which increased their mobility in the softer swelled up matrix when hydrated [51, 83]. This leads to rigid body movements and to a lower strain of fibrils measurable in the axial direction (see also Section 4.2).

Numerically, the effect of hydration was modelled by reducing the collagen stiffness and by using the experimentally found yield strain. The model then predicts a decrease in the apparent fibre mechanical properties including Young’s modulus, yield stress and compressive strength, as seen in experiments. In general we observe a very good agreement between the simulated and experimental results. The model also explains the fibril-to-fibre strain ratio in the elastic region as found experimentally (Figure 6). A decrease of the mineralised collagen fibril stiffness of around 25% led to a reduced apparent fibre stiffness of more than half the value under dry conditions. This decrease in the fibril Young’s modulus agrees with the 15-33% decrease found in molecular dynamics simulations that investigated hydration effects on collagen fibrils deformation mechanisms [51].

### 4.2 Multiscale mechanical behaviour of rehydrated fibres and effect of hydration

Comparing the mineral and fibril strains, and the overall tissue strain to mm-sized compressed bone samples [49], we observe that the influence of hydration differs depending on the constituent. Compared to macroscale samples, fibrils take up 15% less strain and the mineral nearly twice as much (27%). We also see that mineral crystals take 25% less strain in relation to fibrils when directly comparing micrometre (this study) with mm-sized samples [49]. A comparison of the absolute numbers shows that the influence of water on the mineral nanocrystals is 62% higher at the micro- than the macroscale, for the mineralised collagen fibrils the corresponding value is 50%. Interestingly, when we compare the strain ratios with data on mm-sized samples in tension [52], we observe that the decrease of the apparent values is higher in micro-scale samples but the influence of hydration between the constituents is comparable.

Comparing the modelling results for a mineralised collagen fibre under dry and wet conditions, and assuming that all the mineral is located within the fibrils or provided as a mineral impregnation around the fibrils (Section 4.4), we observe that the influence of hydration is 20% higher on the mineral phase than on the mineralised collagen fibrils. One could argue that applying a load directly at the level of an array of mineralised collagen fibrils, one expects a higher strain of the mineral nanocrystals, partly as a result of stress accommodation within the pore spaces between fibres when testing at higher length scales. One explanation for the smaller mineral strains, compared to both the fibril strain and the macroscale (previous paragraph), can be related to the water-mediated structuring of bone apatite [90]. The authors report that the structuring water network facilitates the stacking and orientation of apatite platelets locally. Experimental evidence was given by a higher intensity of the 002 XRD reflection (higher amount of mineral platelets oriented along the main axis of the sample or their preferred orientation, respectively). Compared to X-ray tests under dry conditions [56] we also found that on average the intensity of the 002 signals is higher in wet than in dry samples. Following this rationale, the mineral nanocrystals showed a higher co-alignment in wet samples. Given the high anisotropy of the platelet shape like crystals, in turn, results in a higher overall stiffness in the mineral phase and thus a smaller mineral strain upon loading. Due to the testing at the fibre level, the effect of the water-mediated structuring on apatite is more pronounced in our measured signals. Our data, thus, corroborate the previously reported structural data [90], and outlines the effect on the mechanical behaviour on the micro- and nanometre length scales.

The effect of water on the mineralised collagen fibril strain can be related to a higher mobility of the fibrils within the glue layer of the extrafibrillar matrix as a result of a watery film around the fibrils [51] and a softer swelled up matrix environment [83]. The smaller fibril strains indicate that the interfibrillar gliding due to the watery layer has a more pronounced effect than a reduction of the collagen molecule stiffness (Figure 4). This watery layer might reduce the binding interactions between fibrils and matrix leading to a higher mobility of the fibrils within the extrafibrillar matrix. As a result, a higher amount of energy is dissipated extrafibrillar in the fibril-matrix interactions. The increased mobility might lead to a rigid body movement of the mineralised collagen fibrils within the matrix which in turn results in smaller axial fibril strains since fibrils rather move past each other than being deformed along their longitudinal direction in their uniaxial arrangement [21, 56], which affects the measured SAXS signal.

Comparing the modelling results under dry and wet conditions, the influence of hydration is roughly the same for the strain ratios between mineral nanocrystals and fibrils and those between fibrils and fibre. Assuming that the majority of mineral is located within the fibrils, this points towards three things. First, it indicates that the hydration has a more pronounced influence on the interaction between the mineralised collagen fibrils embedded in the extrafibrillar matrix compared to the intrafibrillar mechanical interaction. Second, it suggests that extrafibrillar energy dissipation mechanisms such as interfibrillar sliding, shear deformation and friction of the matrix glue layer [36, 91–93] are more subject to change during rehydration compared to intrafibrillar mechanisms such as shear strain between the collagen molecules and mineral particles, stress transfer between adjacent mineral particles and intrafibrillar sliding [61, 94, 95]. Third, it supports the suggestion that water is playing an important role in the mediation of mineral-organic matrix interactions [46, 54] within the mineralised collagen fibrils.

### 4.3 Failure mode analysis

For the rehydrated mineralised collagen fibres, the majority of the micropillars failed by mushrooming in combination with either mineralised collagen fibril microbuckling or fracture. In most cases, an interface failure between the mineralised collagen fibril and extrafibrillar matrix was also observed. Furthermore, axial splitting occurred. The dry samples showed the formation of kink bands which led to a localised shear deformation. Axial splitting and localised shear deformation were reported for rehydrated micropillars of ovine bone extracellular matrix [50]. Mushrooming, axial splitting and localised shear deformation were further reported as failure modes for dry tested micropillars of the same material [25]. Thus, in contrast to data reported on rehydrated extracellular matrix bone samples [50], we do not observe kink bands in wet tested mineralised collagen fibres. Considering that we found kink bands in dry tested samples from the same material [56], differences in the fibril orientation might have a more pronounced effect on the failure modes under rehydrated conditions. It can be rationalised by a swelling and cushioning effect due to hydration that leads to an elastic embedding, and translates buckling into well behaved bending. As a result, the formation of kink bands is avoided.

### 4.4 Study limitations

Our rehydration set-up avoided the potential occurrence of bubbles in fully immersed samples (Section 2.1). Preliminary tests were done to quantify the time until HBSS evaporated under ambient conditions while the diffusion state within the micropillar was estimated based on a 1D diffusion calculation (Darcy’s law) [74] (Section 2.1). However, keeping the sample rehydrated before testing by using a pipette for the HBSS refill required special attention in the complex setup of the integrated microindenter near the X-ray tube (Figure 2). An alternative approach would be the use of a climate chamber recently reported for microscale testing where a relative humidity of up to 95% can be reached to simulate physiologically more realistic conditions regarding humidity [96]. This setup was not available for our experiments and needs to be adapted for the combined testing with X-rays. Although being able to simulate the fibre behaviour, our model can only consider a single mineral phase while two are reported to be present with distinct mass fractions. Future work will be directed to this aspect. Using X-rays potentially leads to an irradiation influence on the material [97, 98]. We counteracted this by using short exposure times while keeping the shutter closed in between acquisition times. In addition we used a single-photon counting detector to optimise the time necessary to get a high enough intensity for the X-ray pattern analysis. The mechanical data did not show any effect of the X-ray exposure and the unloading moduli was constant also in the plastic region pointing to an absence of damage (Figure 3).

## 5 Conclusion

We quantified the elasto-plastic micro- and nanoscale behaviour of rehydrated mineralised collagen fibres down to the level of mineral nanocrystals. We used a combined approach of experiments and statistical model to deduce the mineralised collagen fibre behaviour and the effect of hydration. Hydration affected apparent fibre stress more than strain, and we observed an up to three times higher reduction in mechanical properties compared to macroscopic testing which indicates a significant influence of hydration on ultrastructural interfaces in mineralised tissues. Lower stiffness values of the hydrated and swelled excised uniaxial fibril array compared to results at higher scales indicate a loss of reinforcing capacity due to missing surrounding tissue, which is more pronounced than in dry samples. Smaller fibril strains might result from a rigid body movement within the extrafibrillar matrix, and the higher mobility within the fibril-matrix network might lead to a higher amount of energy being dissipated extrafibrillar. The comparable effect of hydration on the compressive strength for bone and mineralised turkey leg tendon indicates that the previously identified difference in mineralisation and interfibrillar surface-to-volume ratio between both mineralised tissues is independent of the rehydration state. However, this also points to the interesting fact that the mobilisation of fibrils follows a similar regime in dry and rehydrated conditions. The decrease in mineral strain was found to be 20% higher than for the fibril strain. Smaller mineral strains under wet conditions can be related to the previously reported water-mediated structuring of bone apatite. The high anisotropy then leads to a higher stiffness in the mineral phase with more co-aligned mineral platelets. During microscale experiments, this effect most likely influences the measured signals more than for macroscale tests. Using a suitable constitutive model allowed us to identify changes at the molecular and fibrillar level as well as the interplay between the constitutive phases that lead to reduced mechanical properties observed experimentally. Our model shows that a decrease of collagen molecule stiffness leads to a reduced fibre yield stress, compressive strength and stiffness while pointing towards differences in the intra- and extrafibrillar phases. The lack of kink bands in the failure mode analysis points towards an elastic embedding of the mineralised collagen fibrils. As for dry conditions, the use of statistically variable micro- and nanoscale mechanical properties were found to be essential for the good agreement with experiments. Our findings close a gap in providing data on the compressive behaviour of single mineralised collagen fibres under rehydrated conditions. Since hydration significantly affects the micro- and nanomechanical properties of mineralised tissues, our results provide invaluable and so far inaccessible information on the understanding of the multiscale elasto-plastic behaviour of bone’s fundamental mechanical unit while pointing towards significant differences to the macroscale.

## 6 Acknowledgements

This research was supported by the Engineering and Physical Sciences Research Council (EPSRC), of the UK (grant number EP/P005756/1) and the European Synchrotron Radiation Facility (ESRF) (proposals ME-1415 and ME-1472). J.S. acknowledges funding by the Swiss National Science Foundation (SNSF) under Ambizione Grant No. 174192. We thank Dr. Richard Carter from the Institute of Photonics and Quantum Sciences at Heriot-Watt University, UK, for organising the access to the laser facilities.

## Appendices

### Determination of strain ratios - Exemplary plots

Figure 8 shows exemplary data to determine the experimental strain ratios between mineralised collagen fibres, mineralised collagen fibrils and mineral nanocrystals.

**Figure 8:**
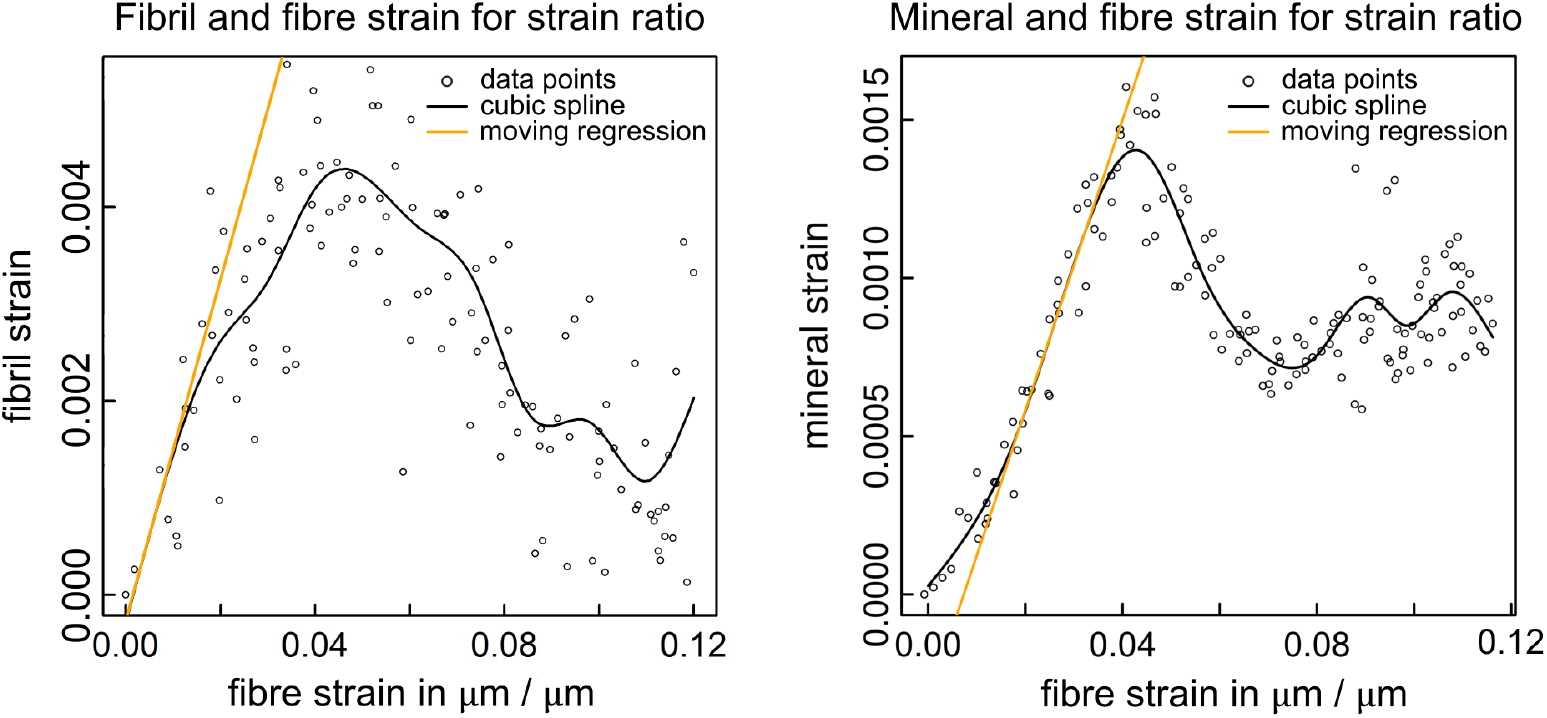
Exemplary data to quantify fibre multiscale strain ratios based on fibril/fibre strain and mineral/fibre strain. A moving regression was used to determine the steepest slope in the elastic region. Section 2.2.

### Overview of model parameters

Table 5 gives an overview of the model input parameters related to composition, values calculated via the two nested shear leg models (elasticity), and those for the plastic material behaviour.

**Table 5:**
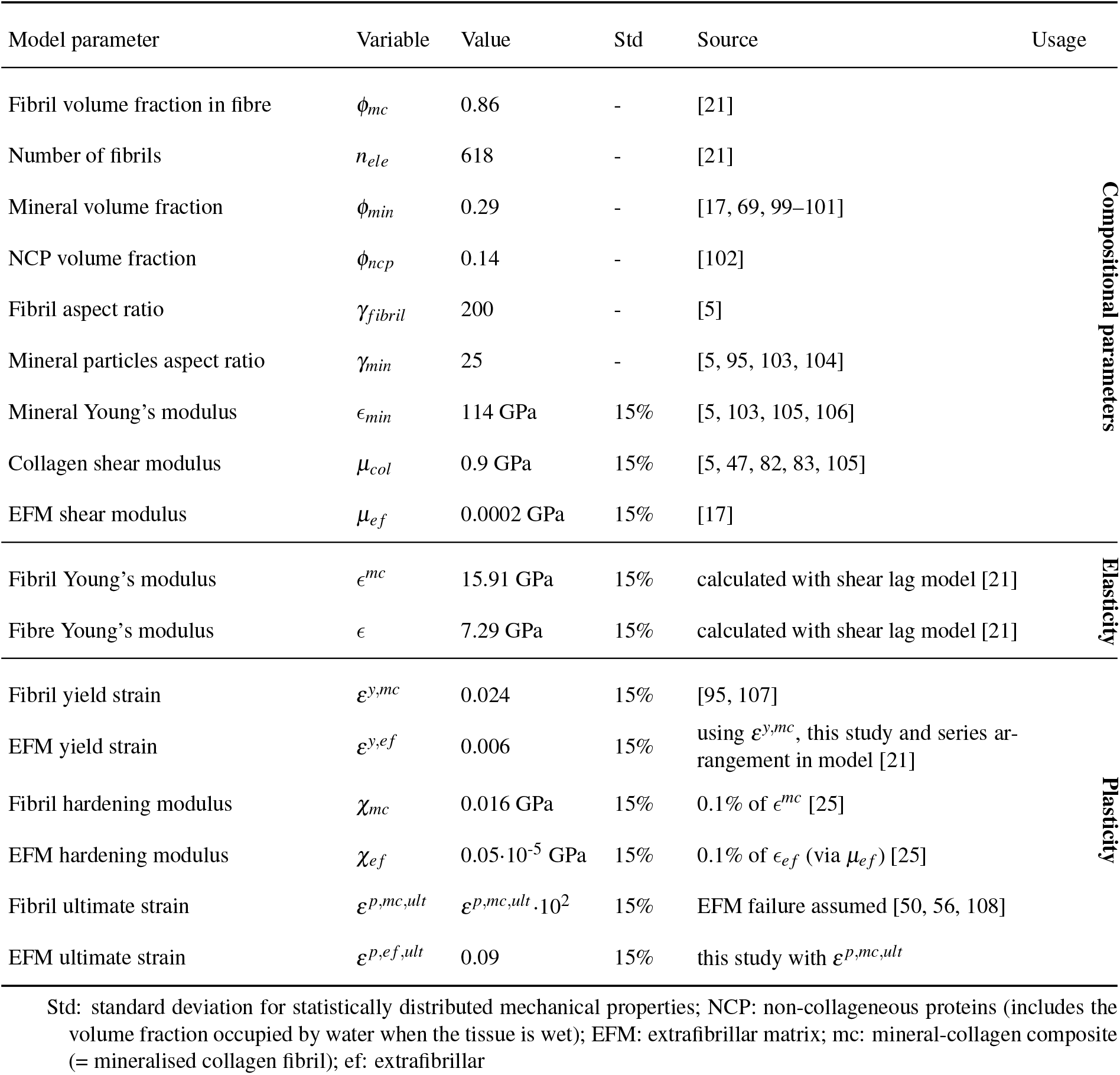
Properties for the quasi-physiologic micro- and nanomechanical behaviour of a mineralised collagen fibre. The table lists the compositional parameters of the fibre (first section) to calculate fibril and fibre elasticity (second section) via two nested shear lag models. All the mineral is assumed to be located inside the fibril/provided as a mineral coating around the fibrils. Plasticity calculations were done based on the elasto-plastic rheological elements (third section). The number of rheological elements equals the number of fibrils. Own experimental values were taken where possible and referenced to the corresponding sections. Additional values were taken from the literature. See [21] for further details.

